# Toxic Effects of Biogenic and Synthesized Silver Nanoparticles on Sea Urchin *Echinometra lucunter* Embryos

**DOI:** 10.1101/2024.04.16.589722

**Authors:** Mariana Bruni, Cristiane Ottoni, Denis Abessa

## Abstract

Due to their broad-spectrum antimicrobial action and ease of synthesis, silver nanoparticles (AgNPs) are one of the most widely used nanomaterials in different industrial and ecological areas. AgNPs are released into marine ecosystems; however, their ecotoxicological effects have been overlooked. In this study, we evaluated the toxic effects of biogenic and synthesized AgNPs on sea urchin *Echinometra lucunter* embryos and compared them to those of AgNO_3_. Fertilized eggs were exposed to five concentrations of the test compounds and a negative control for 48 h under controlled conditions. The EC50-48h of biogenic and synthetic AgNPs and AgNO_3_ were 0.31, 4.095, and 0.01 μg L^-1^, evidencing that AgNPs are less toxic than AgNO_3_, and that synthetic AgNP is less toxic. Toxicity to *E. lucunter* embryos could be explained by the fact that Ag affects DNA replication and induces the formation of pores in the cellular wall, leading to apoptosis.

## 1. Introduction

Nanotechnology is a branch of science and engineering devoted to designing, producing, and using structures, devices, and systems by manipulating atoms and molecules at the nanoscale, i.e., having one or more dimensions on the order of 100 nm or less (Kumar *et al*., 2024). Advances in nanotechnology in recent decades have allowed the large-scale production of manufactured nanomaterials for multiple uses and applications in different fields, such as agriculture, personal care, wastewater treatment, medicine, and packing (Keller *et al*., 2023).

Silver nanoparticles (AgNPs) are emerging nanotechnological compounds applied in different products, mainly due to their recognized antimicrobial properties. Such properties depend on their small size, structure, shape, stability, and environmental physicochemical conditions (Temizel-Sekeryan and Hicks, 2020). According to the Consumer Products Inventory (CPI 2018), from 1814 commercial products containing nanomaterials, 438 consisted of different types of AgNPs produced for a wide variety of uses (Kalantzi *et al*., 2019). The antimicrobial properties of AgNPs are due to many mechanisms of action that affect multiple structures of the target microorganisms and cause their death (Ahmed *et al*., 2024). The total global production of AgNPs was 500 tons in 2009, and it is expected to increase to 900 tons by 2025 (Islam *et al*., 2021). Synthetic AgNPs are generally produced by physical or chemical techniques, whereas the synthesis of biogenic AgNPs has been described as a promising alternative approach because of its low cost and environmentally friendly profile. This process is directly associated with the stabilization and capping of nanoparticles, resulting in reduced toxicity (Santos *et al*., 2024; Silva *et al*., 2022).

The most widely recognized applicability of AgNPs is associated with their antimicrobial activity (Ahmed *et al*., 2024). The mechanism is not completely understood but AgNPs tend to accumulate on biological membranes, forming agglomerates and affecting the cell respiratory chain and phosphate transport, leading to cell death (Nie *et al*., 2023). These nanomaterials have been found in industrial and domestic effluents and can reach the environment. Many effluent treatment systems are incapable to efficiently remove the AgNPs; moreover, many regions around the world do not have appropriate systems to collect and treat the wastewater, being potentially vulnerable to these contaminants (Islam *et al*., 2021). The predicted environmental concentrations of AgNPs were estimated to range between 0.01 and 0.43 μg L^-1^ in aquatic environment (Hlavkova *et al*., 2019).

After environmental exposure, AgNPs interact with the physicochemical components of the environment, and their properties and toxicity may be altered. The ability of AgNPs to interact with biological systems depends on factors such as their size, shape, surface characteristics, and dissolution rates (Rajan *et al*., 2022). Ag ions (including those released from AgNPs) are considered to be persistent and toxic to aquatic organisms (Yang *et al*., 2024). In addition, filter-feeding organisms may have enhanced exposure to NPs, which has implications for their physiology and cell functioning (Sadri and Khoei, 2023). Following the increasing use of AgNPs, studies on the toxicity of these compounds to aquatic organisms have become more frequent in the recent years.

McGillicuddy *et al*. (2017) detected AgNPs toxicity to *Daphnia magna* and the fishes *Danio rerio, Carassius auratus* e *Oncorhynchus mykiss*. The nanomaterial produced effects at the cellular level, affecting embryonic development and causing increased mortality rates. The toxic effects were inversely proportional to NPs size, and the authors attributed this pattern to the higher surface/volume of smaller NPs. The same study observed that while algae and polychaetes accumulated Ag from AgNO_3_ quickly than AgNPs, other invertebrates and fish bioaccumulated the AgNPs.

The aforementioned studies assessed AgNPs toxicity to temperate species, whereas information for polar, tropical, and subtropical species is lacking, despite such being necessary to the understanding of the hazards of NPs to marine species and the establishment of regulations at regional or local levels. In this context, sea urchin embryos have been considered reliable and cost-effective biological models for ecotoxicological studies because they are sensitive and the protocols for toxicity tests are robust and available. Moreover, these organisms are ecologically relevant, have a wide geographical distribution, and are easy to maintain in the laboratory (Environment Canada, 1992). In Brazil, *Echinometra lucunter* has been used in ecotoxicological studies. This organism is abundant along the Brazilian coast and is considered to be a suitable test species (Nascimento *et al*., 2002).

This study aimed to evaluate and compare the potential toxicity of synthetic and biogenic compounds in the embryonic development of *E. lucunter*. This also provides the first results regarding these AgNPs in the marine species of Brazil.

## 2. Materials and Methods

### 2.1 Ag Nanoparticles

*Penicillium citrinum* IBCLP11 was grown on PDA for seven days, and colonies were used for biogenic AgNP synthesis. Briefly, the formation of biogenic AgNPs_IBCLP11_ was initially confirmed by UV-visible spectrophotometry, which demonstrated a plasmon band resonance at 427 nm. The AgNPs particle size was determined by transmission electron microscopy (TEM), which showed an average diameter of 57.5 ± 4 nm. The polydispersity index (PDI) and zeta potential (Pζ) were 0.207 and -41.91, respectively (Aguiar *et al*., 2024). Synthetic AgNPs were synthesized by chemical reduction using polyvinyl pyrrolidone (PVP), their Ag content was 0.2 mg mL^-1^ and average diameter of 17 nm.

### 2.2 Toxicity tests

The toxicity tests followed the protocols established for embryos of *E. lucunter* and neotropical echinoderms (Nascimento *et al*., 2002; ABNT 2012), which are based on international guidelines (Environment Canada, 1992). Basically, the tests consisted of exposing fertilized eggs of *E. lucunter* to solutions of the substances tested, considering five concentrations ranging between 0.01 and 100 μg L^-1^, under controlled physical-chemical conditions. Four replicates were prepared for each test solution and the negative control (filtered clean seawater). After approximately 48h, the tests were completed and we observed successful embryonic development and the presence of morphological abnormalities in the larvae. The results obtained for each substance were analyzed by one-way Analysis of Variance (ANOVA) followed by Dunnett’s test to determine the Lowest Observed Effect Concentrations (LOECs) and the No Observed Effect Concentrations (NOECs). Additionally, the inhibition concentrations in 50% embryos after 48h (IC_50-48h_) were estimated by non-linear regression. During the experiments, the physicochemical parameters of all solutions were measured at the beginning and end of the tests.

### 2.3. Chemical Analysis

The concentrations of Ag in the stock solutions of AgNO_3_ and biogenic and synthetic AgNPs were analyzed using Flame Atomic Adsorption Spectrometry (FAAS). Aliquots (0.75 mg L^-1^) of each substance were prepared for analysis. The calibration curve was obtained after the analysis of five Ag solutions (1, 0.75, 0.5, 0.25, and 0.1 mg L^-1^), and the calibration curve obtained showed an r^2^ value greater than 0.99, with the equation y = 0.895x – 0.0005, which was used to adjust the nominal concentrations to the real ones used in the toxicity test. All solutions were prepared with deionized water and read in the spectrometer at a 328.1 nm wavelength. The quantification limit was 0.6 μg L^-1^ and detection limit was 0.18 μg L^-1^. All reagents used were grade standards.

## 3. Results

### 3.1 Chemical Analysis Results

The Ag concentrations in the stock solutions were 15.4 mg L^-1^ in the AgNO_3_ solution, and 0.57 and 1.89 mg L^-1^ in the biogenic and synthetic AgNPs, respectively.

### 3.2 Ecotoxicity Results

In the experiments, the physicochemical parameters remained within the ranges established for the test (ABNT, 2012), as shown in Supplementary Material. Salinity remained constant, and slight fluctuations in pH and dissolved oxygen were within acceptable ranges. For the biogenic AgNPs_IBCLP11_ (Figure 1A), the mean normal embryonic development rate in the controls was 95±2.9% The LOEC was estimated as 0.37 μg L^-1^, while the NOEC was estimated as 0.22 μg L^-1^ (Table 1). At higher concentrations, 100% of embryos were affected. The IC_50-48h_ of the biogenic AgNPs_IBCLP11_ was calculated as 0.31 μg L^-1^ (0.28 – 0.35). The synthetic AgNPs induced significant effects (p<0.05) from the 0.25 μg L^-1^ concentration, presenting a NOEC of 0.025 μg L^-1^ (Figure 1B). The IC_50-48h_ was estimated as 4.095 μg L^-1^ (0.7987 - 20.99). In the test with AgNO_3_, the LOEC was 0.002 μg L^-1^, and the NOEC was 0.0002 μg L^-1^ (Figure 1C). Embryonic development was not observed at the highest concentrations there was no. For this compound, the IC_50-48h_ was calculated as 0.01 μg L^-1^ (0.004 – 0.03).

**Table 1.** Summary of the results obtained in the toxicity tests with the biogenic AgNP (AgNP_IBCLP11_), synthetic AgNPs (AgNPs sin) and AgNO_3_, showing the respective Lowest Observed Effect Concentrations (LOECs), No Observed Effect Concentrations (NOECs), and inhibition concentrations to 50% embryos of Echinometra lucunter after 48h (IC50-48h).

| Substance | NOEC | LOEC | $\text{IC}_{50}$ |
| --- | --- | --- | --- |
| $\text{AgNPs}_{\text{IBCLP11}}$ | 0.22 $\mu\text{g L}^{-1}$ | 0.37 $\mu\text{g L}^{-1}$ | 0.31 $\mu\text{g L}^{-1}$ (0.28- 0.35) |
| $\text{AgNPs}_{\text{sin}}$ | 0.025 $\mu\text{g L}^{-1}$ | 0.25 $\mu\text{g L}^{-1}$ | 4.095 $\mu\text{g L}^{-1}$ (0.7987 - 20.99) |
| $\text{AgNO}_3$ | 0.0002 $\mu\text{g L}^{-1}$ | 0.002 $\mu\text{g L}^{-1}$ | 0.01 $\mu\text{g L}^{-1}$ (0.004 -0.027) |

**Figure 1.**
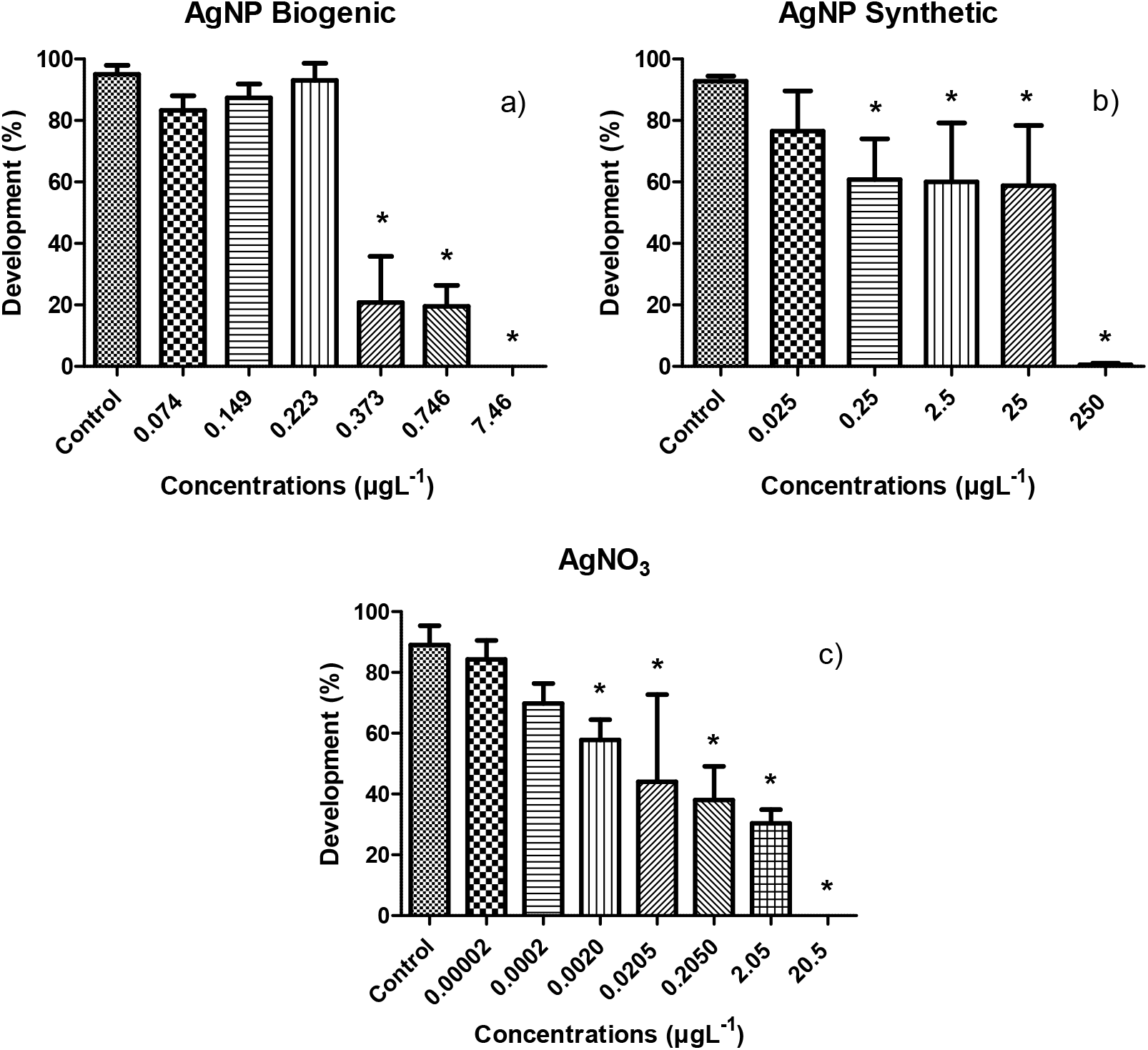
Normal embryonic development rates of *Echinometra lucunter* exposed to different concentrations of a) Biogenic AgNPs_IBCLP11_, b) Synthetic AgNPs, and c) silver nitrate (AgNO_3_). Asterísks (*) indicate significant differences to the respective controls (p<0.05).

## 4 Discussion

Assessing the toxicity of different types of AgNPs is important for developing new products containing such compounds. In this study, adverse effects in *E. lucunter* embryos occurred at lower concentrations of all the substances tested. Significant adverse effects were observed in embryos exposed to 0.25 μg L^-1^ synthetic AgNP, 0.002 μg L^-1^ AgNO_3_, and 0.373 μg L^-1^ biogenic AgNPs_IBCLP11_, indicating the high sensitivity of these young life forms to AgNPs and their precursor salt AgNO_3_. AgNPs were much less toxic to *E. lucunter* embryos than AgNO_3_, and biogenic AgNPs_IBCLP11_ was less toxic than synthetic AgNP.

Our results suggest that *E. lucunter* embryos are more affected by synthetic AgNPs and their precursor metals than their temperate counterparts, following the results of Siller et al. (2013). Embryos of *Paracentrotus lividus* showed delayed development, and from the concentration of 0.3 mg L^-1^, the AgNP become less toxic than the free Ag (0.003 mg L^-1^). Buric et al. (2015) exposed fertilized eggs of three species of sea urchins, *Arbacia lixula, Paracentrotus lividus*, and *Sphaerechinus granularis*, to different concentrations of AgNP and AgNO_3_, ranging from 1 to 100 μg L^-1^, and observed adverse effects at concentrations of 1-10 μg L^-1^. In that case, *A. lixula* was the most sensitive species; thus, the authors concluded that the effects of AgNPs were species-specific. Although *E. lucunter* is from a different environment, when we compared the results obtained for this tropical species with those of the temperate species, tropical organisms were more sensitive. Another factor that might interfere with the toxicity results is the synthesis method. The method used by Buric et al. (2015) to synthesize AgNPs was based on reduction with sodium citrate, and the method used in the present study was reduction with PVP.

Information obtained from freshwater organisms is also reported in the literature. Zhao and Wang (2011) compared the toxicity of AgNP and AgNO_3_ to *Daphnia magna* and observed that AgNO_3_ was more toxic (LC50 = 2.51 μg L^-1^), while the AgNP did not induce significant toxicity. Ribeiro et al. (2014) also compared the toxicities of AgNPs and AgNO_3_ to *D. magna* after 24 and 48h, and observed that the AgNPs were ten times less toxic than the AgNO_3_, similarly to our results. In these cases, Ag dissolution seems to be the main factor influencing toxicity, and the lower toxicity of AgNPs is likely due to the fact that NPs control the release of reactive Ag into the water column (Ribeiro et al., 2014). On the other hand, Asharani et al. (2008) observed that AgNPs were more toxic than AgNO_3_ to the embryonic development of *Danio rerio*, possibly because AgNP can penetrate into the chorionic membrane and interact with the embryo tissues.

The toxicity of AgNPs is mainly caused by Ag ions, which have a tendency to bioconcentrate in organisms, as they are compatible with molecules transported through the cell membrane, such as thiol groups of enzymes and proteins. Such interactions may be responsible for the influx of ions into cells, causing organelle damage and blocking cell division (Fabrega et al., 2010). The free ions of Ag compete with sodium for the connection sites of the enzyme Na^+^/K^+^-ATPase, which is important for capturing and transporting Na^+^ and Cl^-^ from water to invertebrate extracellular fluids, causing insufficient osmoregulation and death (Ribeiro et al., 2014). Ag ions may interact with cellular components, altering metabolic routes and genetic material (Nie et al., 2023), interfering with the immune system, and causing enzyme inactivation and depolarization of the mitochondrial membrane (Vali et al., 2022).

In turn, AgNPs present multiple mechanisms of action that cause toxicity, including those caused by Ag ions, as reported above. The particles may penetrate the cell membrane and accumulate in their internal layer, producing membrane destabilization and damage that alter permeability. Such effects lead to leaching of cellular content and cell death. In addition, AgNPs may bind to cellular molecules containing sulfur and phosphorus (such as DNA and proteins), leading to alterations in their structure and function, and inducing oxidative stress. The combination of these multiple effects induces apoptosis, cytotoxicity, and genotoxicity, which could explain the toxic effects in the embryos of *E. lucunter*, since the inhibition of cell division possibly prevented the development of embryos.

## 5 Conclusion

Embryos of *E. lucunter* exhibited higher sensitivity to AgNO_3_ than to AgNPs, as evidenced by their respective LOECs and IC_50-48h_ values. However, the effects of the synthetic AgNPs start slightly early, with a LOEC of 0.25μg L^-1^ against 0.37 μg L^-1^ of biogenic AgNPs_IBCLP11_.

## 6 Aknowledgments

This research was funded by the São Paulo Research Foundation (FAPESP; grants #2020/03004-0; #2020/12867-2). Mariana Bruni thanks FAPESP for the scholarships (grant #2018/25379-6). The authors are grateful to Prof. Dr. Emília Lima (UFG) for providing the compounds, and Drs. Fernando Perina (UA) and Caio Ribeiro (UNESP) for their contributions.

