## Supplementary Material for "Toxic Effects of Biogenic and Synthesized Silver Nanoparticles on Sea Urchin *Echinometra lucunter* Embryos"

**Table 1.** Embryo development of *Echinometra lucunter* exposed to biogenic $AgNPs_{IBCLP11}$. The hifen (-) means the absence of replicate.

| Control | 0.074 µgL$^{-1}$ | 0.149 µgL$^{-1}$ | 0.223 µgL$^{-1}$ | 0.373 µgL$^{-1}$ | 0.746 µgL$^{-1}$ | 7.46 µgL$^{-1}$ |
|---|---|---|---|---|---|---|
| 92 | 78 | 92 | 85 | 37 | 11 | 0 |
| 94 | 81 | 87 | 94 | 9 | 17 | 0 |
| 95 | 89 | - | 95 | 30 | 24 | 0 |
| 99 | 85 | 83 | 98 | 7 | 26 | 0 |

**Table 2.** Physical-chemical parameters of biogenic $AgNPs_{IBCLP11}$ during the exposition time.

| | pH | | DO (mg L$^{-1}$) | | salinity | |
|---|---|---|---|---|---|---|
| **Concentrations** | **inicial** | **final** | **inicial** | **final** | **inicial** | **final** |
| 0.074 µg L$^{-1}$ | 7.82 | 7.87 | 6.69 | 6.42 | 34 | 34 |
| 0.149 µg L$^{-1}$ | 7.58 | 7.9 | 6.6 | 6.41 | 34 | 34 |
| 0.223 µg L$^{-1}$ | 7.33 | 7.77 | 6.75 | 6.52 | 34 | 34 |
| 0.373 µg L$^{-1}$ | 7.72 | 7.9 | 6.66 | 6.45 | 34 | 34 |
| 0.746 µg L$^{-1}$ | 7.73 | 7.63 | 6.68 | 6.66 | 34 | 34 |
| 7.46 µg L$^{-1}$ | 7.18 | 7.74 | 6.91 | 6.38 | 34 | 34 |
| Control | 7.99 | 6.73 | 7.92 | 6.88 | 34 | 34 |

**Table 3.** Embryo development of *Echinometra lucunter* exposed to synthetic AgNPs.

| Control | 0.025 µgL$^{-1}$ | 0.25 µgL$^{-1}$ | 2.5 µgL$^{-1}$ | 25 µgL$^{-1}$ | 250 µgL$^{-1}$ |
|---|---|---|---|---|---|
| 93 | 83 | 44 | 59 | 56 | 0 |
| 95 | 81 | 64 | 83 | 32 | 1 |
| 91 | 85 | 76 | 36 | 73 | 1 |
| 92 | 57 | 59 | 62 | 74 | 0 |

**Table 4.** Physical-chemical parameters of synthetic AgNPs during the exposition time.

| | pH | | DO (mgL$^{-1}$) | | salinity | |
|---|---|---|---|---|---|---|
| **Concentrations** | **inicial** | **final** | **inicial** | **final** | **inicial** | **final** |
| 0.025 µg L$^{-1}$ | 7.18 | 7.74 | 6.75 | 6.06 | 34 | 34 |
| 0.25 µg L$^{-1}$ | 7.2 | 7.72 | 6.79 | 6.44 | 34 | 34 |
| 2.5 µg L$^{-1}$ | 7.21 | 7.85 | 6.84 | 6.35 | 34 | 34 |
| 25 µg L$^{-1}$ | 7.22 | 7.92 | 6.66 | 6.60 | 34 | 34 |
| 250 µg L$^{-1}$ | 7.32 | 7.24 | 6.76 | 6.88 | 34 | 34 |
| Control | 7.99 | 6.73 | 7.92 | 6.88 | 34 | 34 |

**Table 5.** Embryo development of *Echinometra lucunter* exposed to synthetic $AgNO_3$.

| Control | 0.00002 µgL$^{-1}$ | 0.0002 µgL$^{-1}$ | 0.002 µgL$^{-1}$ | 0.0205 µgL$^{-1}$ | 0.205 µgL$^{-1}$ | 2.05 µgL$^{-1}$ | 20.5 µgL$^{-1}$ |
|---|---|---|---|---|---|---|---|
| 94 | 87 | 60 | 56 | 63 | 49 | 30 | 0 |
| 80 | 75 | 71 | 51 | 11 | 43 | 57 | 0 |
| 93 | 86 | 74 | 57 | 0 | 37 | 26 | 0 |
| 89 | 89 | 74 | 67 | 58 | 23 | 35 | 0 |

**Table 6.** Physical-chemical parameters of synthetic $AgNO_3$ during the exposition time.

| | pH | | DO (mgL$^{-1}$) | | salinity | |
|---|---|---|---|---|---|---|
| **Concentrations** | **inicial** | **final** | **inicial** | **final** | **inicial** | **final** |
| 0.00002 µg L$^{-1}$ | 7.82 | 7.68 | 6.72 | 6.81 | 34 | 34 |
| 0.0002 µg L$^{-1}$ | 7.75 | 7.84 | 6.37 | 6.87 | 34 | 34 |
| 0.002 µg L$^{-1}$ | 7.75 | 7.27 | 6.68 | 6.59 | 34 | 34 |
| 0.0205 µg L$^{-1}$ | 7.83 | 7.34 | 6.37 | 6.60 | 34 | 34 |
| 0.205 µg L$^{-1}$ | 7.76 | 7.55 | 6.67 | 6.66 | 34 | 34 |
| 2.05 µg L$^{-1}$ | 7.78 | 7.92 | 6.58 | 6.85 | 34 | 34 |
| 20.5 µg L$^{-1}$ | 7.86 | 7.76 | 6.59 | 6.73 | 34 | 34 |
| Control | 7.99 | 6.73 | 7.92 | 6.88 | 34 | 34 |
